# Understanding Shape and Residual Stress Dynamics in Rod-Like Plant Organs

**DOI:** 10.1101/2025.08.25.672092

**Authors:** Amir Porat

## Abstract

Residual stresses are common in rod-like plant organs such as roots and shoots, arising from mechanical incompatibilities between tissues with differing intrinsic lengths. Although mechanical regulation of cell wall growth is well established, the role of internally generated stresses in organ-scale morphogenesis remains poorly understood. Here, we introduce a tractable theoretical framework that couples structural residual stresses to growth-driven shape dynamics. We represent rod-like organs as bundles of morphoelastic rods connected in parallel, each corresponding to a distinct concentric tissue layer. Assuming elastic strain-driven growth and rod-like symmetry, the resulting mechanical constraints yield transparent analytical relationships linking tissue heterogeneity to macroscopic observables such as axial strain rates, bending dynamics, and residual stress distributions. Using a minimal two-layer model representing the epidermis and inner tissues, we show how tissue incompatibilities can generate phenomena including effective autotropism, mechanical memory, and discontinuous growth dynamics following cell division. Overall, our framework demonstrates how residual stresses may contribute to plant morphogenesis and provides a foundation for future investigations of mechano-chemical growth control in plants.

## Introduction

Residual stresses are widespread in plants and play a key role in function, development, and mechanical stability. However, how they influence macroscopic shape dynamics remains poorly understood (1–7). Residual stresses, defined as stresses that persist within tissues in the absence of external forces, can arise from internal turgor pressure which maintains cell wall tension (3, 8), or from structural incompatibilities between tissues with different intrinsic lengths. Such incompatibilities may result from differential growth rates (9) or tissue shrinkage, as observed in tendrils and reaction wood (10–12) (Fig. 1A). Since plant growth is coupled to its elastic state (13, 14), understanding the development and distribution of residual stresses is essential for studying plant morphogenesis. However, as mechanical fields such as stress and strain are notoriously difficult to measure experimentally, investigations into their developmental role often rely on physics-based modeling. Within this interdisciplinary effort, non-lignified rod-like organs such as roots and shoots provide an attractive model system: they exhibit strong axial growth symmetry that preserves their slender geometry, while also enabling controlled experimental perturbations through tropic bending responses to external directional stimuli (15). To explore the interplay between growth and elasticity in these systems, two major classes of theoretical models are often applied. First are morphoelastic rods, which describe organ-scale morphogenesis through the spatiotemporal dynamics of a growing one-dimensional centerline, often coupling local growth to morphogens such as auxin (9, 16–18). While successful in reproducing certain movements such as tropisms, circumnutations, and mechanical interactions with the environment, these models generally prescribe growth fields and treat mechanical stresses as downstream consequences. As a result, they either neglect elastic deformations as a whole (18–22), ignore possible feedback from tissue mechanics on growth (23–25), or average elastic fields over cross-sections (16).

**Fig. 1.**
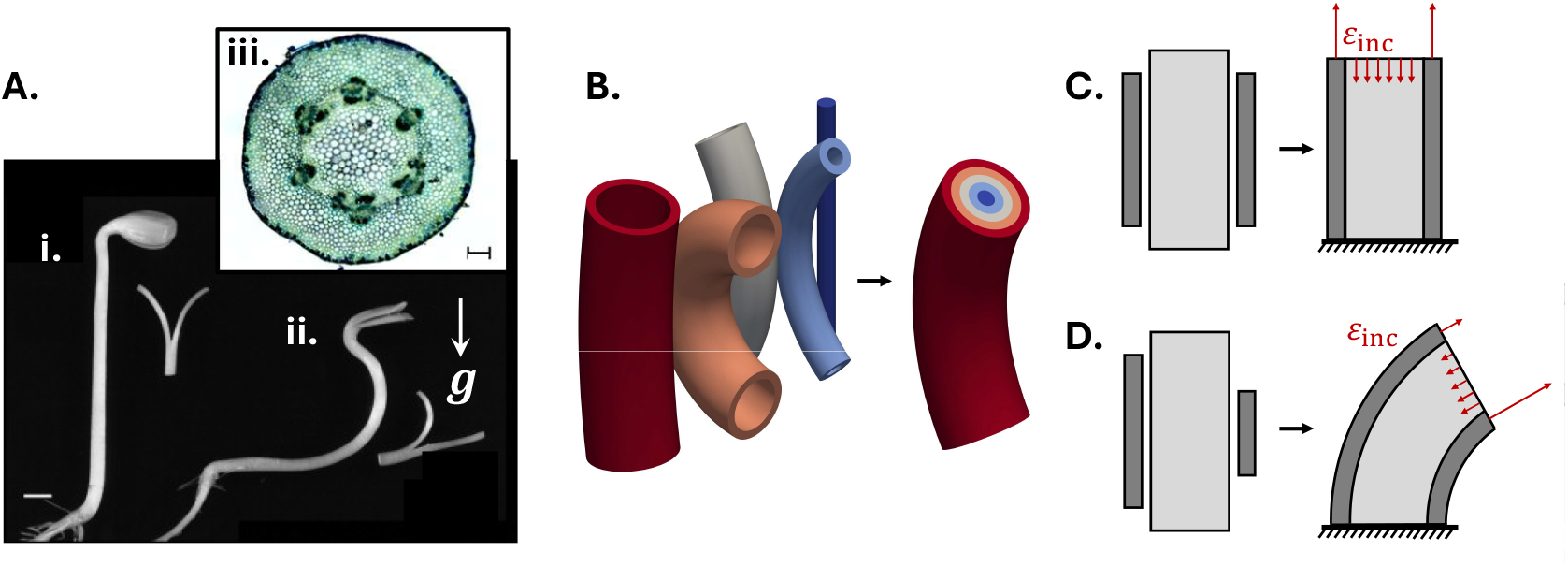
A. Biological motivation. i.-ii. Photograph of 4-day-old etiolated sunflower seedlings and hypocotyl segments that were split longitudinally and placed for 1h into water. i. maintained vertically or ii. placed for 3h in horizontal position to induce gravitropic upward bending. The split halves curved outward, reflecting non-trivial residual stresses. ***g*** is the direction of gravity, scale bar=5mm. Taken from (1). iii. A cross section of a sunflower hypocotyl, showing concentric arrangement of various tissues. Image courtesy of Ahron Kempinski, scale bar 0.2mm. **B. Bundle of concentric shells**. To model the kinematics of an organ with internal residual stress, we use *N* connected uniform concentric cylindrical shells which grow together and constrain each other elastically. **C-D. Elastic strain due to structural incompatibility**, *ε*_inc_. Each illustration depicts a section of a bundle composed of two concentric shells, as in (B). **C**. Typical residual stress profile in rod-like plant organs, where the epidermis is under tension and the inner tissue is relatively compressed. This profile may also arise from differential stiffness (see Eq. 58 in the SI). **D**. Differential axial growth, superimposed on the residual strain profile in (C), produces bending and results in an asymmetric distribution of residual strain.

The second type of models work on the cellular-scale, incorporating experimentally-inspired cell architectures, solve their mechanical equilibrium under turgor, and integrate growth-driven shape changes based on assumed growth laws (26–29). These detailed models are able to provide a more complete description of the interplay between genetics, cell wall growth, and mechanics. Nevertheless, the computational cost and high parameter complexity of these models make systematic exploration of long-timescale morphodynamics difficult.

Consequently, the potential role of mechanical incompatibilities between tissues in shaping organ-scale morphodynamics remains largely unresolved. To address this gap, we introduce an intermediate-scale modeling framework, in which rod-like organs are represented as bundles of morphoelastic, concentric cylindrical shells, reflecting the longitudinal organization of cell files into distinct tissues in roots and shoots (Fig. 1A,B). Assuming quasi-static growth proportional to local elastic strain, with each shell possessing distinct elastic and growth properties, we compute mechanical equilibrium a priori and derive ODEs governing axial growth and curvature, enabling both numerical integration and analytical insight into the coupling between growth and elasticity in plants.

As a proof of concept, we focus on a minimal two-layer model representing the epidermis and inner tissues, inspired by the “epidermal growth control” hypothesis which proposes that the epidermis mechanically constrains inner tissue expansion (1, 30). We use this minimal model to investigate several case studies, including elastic relaxation following cell division, a novel mechanical model for autotropism and memory effects in bending motions, and a framework for exploring mechanochemical feedback of morphogens on stationary axial growth profiles.

Our theoretical framework and accompanying case studies highlight the substantial dynamical role that residual stresses may play in plant development, while providing a transparent mathematical framework for their investigation in future work.

### A single morphoelastic rod

We begin by mathematically describing the shape kinematics of an axially growing morphoelastic rod which is homogeneous in all elastic and growth related properties. Building on previous models (9, 16, 25), the rod geometry is depicted using its centerline ***r***(*s, t*), parametrized by its arc-length *s* and time *t*, as well as a local orthonormal material frame {**ê**_1_(*s, t*), **ê**_2_(*s, t*), **ê**_3_(*s, t*)} (17, 31) (see table of parameters used in the supplementary information (SI)). Material points around the centerline are described using a vector on the local cross section ***x*** = *x*_1_**ê**_1_ + *x*_2_**ê**_2_, and the shape of the cross section is assumed to be planar and constant (see Fig. 2A). Assuming the rod is twistless and shearless allows us to identify the tangent to the centerline 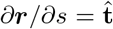 as a material frame vector 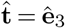 and define the curvature or Darboux vector as 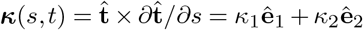. The shape kinematics is then fully described by the axial-axial component of the deformation rate tensor **d**, which we term the axial strain rate and denote using 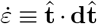 (see SI for derivation). Denoting co-rotating material time derivatives using 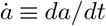 such that 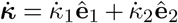, and assuming small curvatures such that |***κ*** × ***x***| ≪ 1 for all points on the cross section, we find that the axial stain rate is linear within the cross section:

**Fig. 2.**
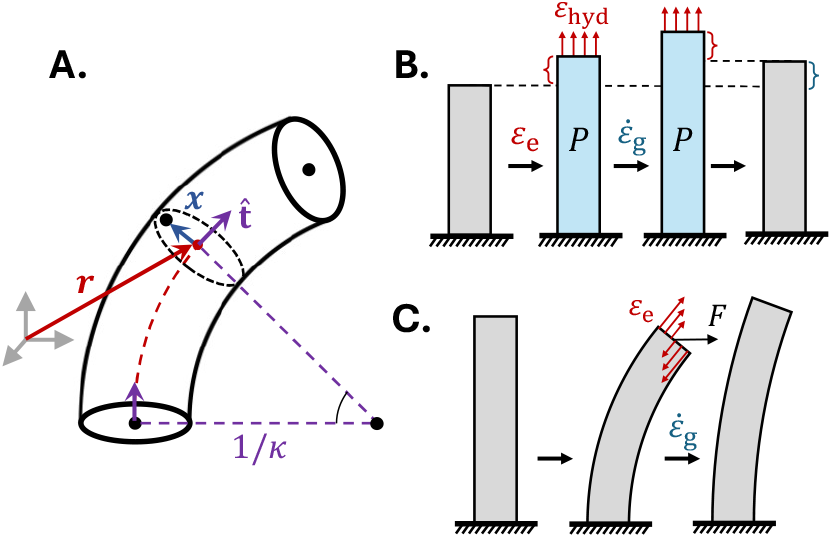
A single morphoelastic rod. A. Geometry. A morphoelastic rod is defined by its moving centerline ***r***(*s, t*) defined in arc length *s* and time *t*, as well as a local material frame {**ê**_1_(*s, t*), **ê**_2_(*s, t*), **ê**_3_(*s, t*)} (not shown). We define the cross section of the rod *A* as the set of material points perpendicular to the tangent to the centerline 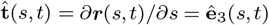, effectively assuming a shearless rod, and span the cross section using the vector ***x***(*s, t*) = *x*_1_**ê**_1_(*s, t*) + *x*_2_**ê**_2_(*s, t*). The rotation of the tangent along the centerline is captured by the curvature vector ***κ***(*s, t*) = *κ*_1_**ê**_1_ + *κ*_2_**ê**_2_, as represented here by an organ with constant curvature |***κ***| = *κ* and radius of curvature 1*/κ*. **B. Morphoelasticity**. Elastic strain (*ε*e) in response to internal hydrostatic turgor pressure (*P*) elongates the tissue, while growth 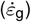 increases the intrinsic length of the organ - both contributing to the total observed deformation. **C. Intrinsic curvature dynamics**. Axial strain-based growth 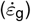 in the presence of elastic bending deformations (*ε*_e_) induces changes in the intrinsic curvature. Here, bending results from the moment caused by an external force *F* .

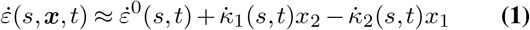

where 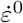 is the strain rate of the centerline, and the linear terms are related to the curvature rate 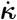. In the following, we use this linearity to completely separate the dynamics of curvature and average axial growth. We now assume that external forces elastically deform the intrinsic shape of the rod. For example, when external moments are applied, the intrinsic curvature ***κ***^0^ may differ from the current actual curvature ***κ***, implying elastic bending strains. For simplicity, we restrict the discussion to bending and stretching deformations and assume no elastic shear or twist, thus considering only axial stresses and strains. The Cauchy stress tensor can then be written as 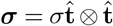, and the distribution of axial elastic strain *ε*_e_ as (32, 33):

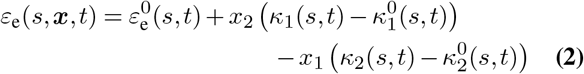

where 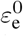 is the average elastic axial strain. Taking a coarse-grained approach (3, 34, 35), we assume linear elasticity such that:

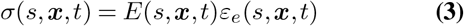

where *E* is an effective Young’s modulus describing tissue-scale elastic behavior. Unless mentioned otherwise, we constrain *E* to be time-independent and axisymmetric such that *E*(*s*, ***x***, *t*) = *E*(*S*_0_, |***x***|), where *S*_0_ is a material coordinate (see SI for details), and assume its first moment on each cross section vanishes.

#### Strain-based growth leads to stress relaxation

Due to the internal hydrostatic turgor pressure in living plant cells, the surrounding stiff cell walls are typically under tension. Growth proceeds via creep-like deformations driven by this wall tension. Because growth occurs on timescales much slower than viscoelastic relaxation, we can assume quasistatic shape dynamics, i.e. the morphoelastic rod remains in mechanical equilibrium at all times (17). The axial strain rate can then be decomposed additively to the axial relative growth rate 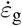 and the quasi-static variations of axial elastic strain 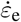 via 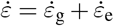 (17, 36) (see Fig. 2B). In plants, growth is commonly described by the Lockhart model, which relates the anisotropic elongation of cell walls to the internal turgor pressure *P* (13, 35, 37, 38). Following recent generalizations of this model (17, 39–41), we adopt a strain-based growth formulation in which the axial growth rate is proportional to the axial elastic strain in the cell wall. Specifically, we use the relation:

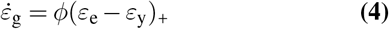

where *ϕ* is the effective extensibility (with units 1/time), *ε*_y_ is the yield strain, and (*x*)_+_ ≡ *x*Θ(*x*) where Θ is the Heaviside function. The total axial strain rate can then be written as:

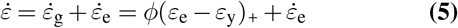

If the strain rate 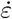 is known, Eq. 5 represents a differential equation for *ε*_e_. For example, a uniform morphoelastic organ growing axially under geometrical confinement acts as a loaded column with constant length and 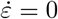, simplifying the resulting ODE. Then, assuming no arc-length dependence, *ε*_e_(*t* = 0) = *ε*_*e*,0_ *> ε*_y_, and no buckling, Eq. 5 gives *ε*_*e*_(*t*) = *ε*_y_ + (*ε*_*e*,0_ − *ε*_y_)*e*^*−ϕt*^, such that the elastic strain decays exponentially to its yield threshold (42). As for the dynamics of intrinsic curvature ***κ***^0^, in the SI we show that when the rod grows everywhere (*ε*_e_ *> ε*_y_), Eqs. 1-5 give:

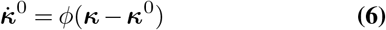

such that the intrinsic curvature ***κ***^0^ relaxes to its actual curvature ***κ*** in the presence of external forces (43) (see Fig. 2C). In general, elastic bending may produce compressive elastic strains that go below the yield threshold (*ε*_e_ ≤ *ε*_y_), causing growth to cease only on one side of the organ, as described in Eq. 2. Since the resulting growth pattern breaks the planarity of the intrinsic cross-section, the ensuing bending dynamics become more complex and lie outside the scope of this paper (for a full continuous treatment see, for example, (9)).

Equation 6 describes the passive response to elastic bending deformation, however, plant rod-like organs often actively vary their intrinsic curvature, as exemplified by tropic and nastic movements (12, 16, 18, 30, 44, 45). Within the strain-based growth model, such movements can arise from spatial variations in growth parameters (*ϕ, ε*_y_), stiffness (*E*), or internal turgor pressure (*P*) across the rod cross-section. To illustrate this, we assume the shape dynamics of the centerline are planar with *κ*_1_ = 0, and that *ϕ, ε*_y_, *E* and *P* vary linearly in the plane of bending (*x*_1_, *s*). By denoting *da/dx*_1_ ≡ *a*^*′*^, this translates to *E*(*x*_1_, *t*) = *E*^0^(*t*) + *x*_1_*E*^*′*^(*t*), *P* (*x*_1_, *t*) = *P* ^0^(*t*)+*x*_1_*P*^*′*^(*t*), 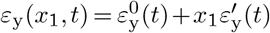 , and *ϕ*(*x*_1_, *t*) = *ϕ*^0^(*t*) + *x*_1_*ϕ*^*′*^(*t*). Then, assuming the variations are small, we obtain two expressions to the first order in the spatial gradients (see SI). First, a non-trivial bending strain Δ*κ*_hyd_ arising from internal hydrostatic turgor pressure in the presence of transverse gradients in either turgor or stiffness:

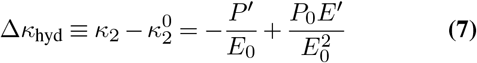

Second, an updated dynamics of intrinsic curvature:

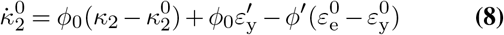

where the first term on the right hand side reproduces Eq. 6, which in the absence of external forces can be attributed to Δ*κ*_hyd_ as shown in Eq. 7. Given an observed growth-driven bending motion, distinguishing experimentally between the bending mechanisms described by Eqs. 7 and 8 remains an open challenge. Under osmotic treatments, however, the magnitude of Δ*κ*_hyd_ is expected to vary, leading to observable changes in curvature. This could provide a means to discriminate between bending driven by gradients in growth parameters and that arising from variations in turgor or stiffness.

Conceptually, Eqs. 5–8 show that in a homogeneous morphoelastic rod, which does not contain residual stresses, a strain-driven growth law leads to exponential relaxation of the curvature and axial strain rates toward steady-state values. As we show next, once the rod becomes inhomogeneous and distinct tissues are allowed to mechanically constrain one another during growth, these steady-state values naturally depend on the heterogeneity within the organ.

### A bundle of N concentric morphoelastic cylindrical shells

#### Model assumptions

We now model the rod-like growing organ as a bundle of *N* connected, concentric cylindrical shells, each representing a distinct tissue layer (see Fig. 1B). While elastic and growth-related parameters may vary between shells, we assume that within each shell these properties are uniform, and that axial growth is compatible and follows Eqs.5,6. Although the bundle grows freely - without external loads - the shells are mechanically coupled in parallel. This simulates symplastic growth, in which connected plant cells expand cohesively, without slippage between cells nor tissue layers. For each shell *i* we therefore decompose the elastic strain (or stress) into two additive contributions:

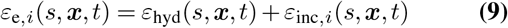

where *ε*_hyd_(*s*, ***x***, *t*) are reaction strain to the internal hydrostatic turgor pressure and *ε*_inc_(*s*, ***x***, *t*) are residual strains caused by structural incompatibility (see Fig. 1C,D). Under the assumptions of linear elasticity and a planar cross-section, we show in the SI that for constant turgor pressure *P* , *ε*_hyd_ is uniform across the cross-section and equals *P/* ⟨*E*⟩ , where ⟨*E*⟩ denotes the average (bulk) stiffness of the cross-section under pressurization. To continue, we make the following simplifying assumptions: (i) the curvature of each shell is uniform along its length; (ii) the problem is two dimensional: the centerline and plane of curvature are on the (*x*_1_, *s*) plane (curvature vectors are parallel or anti-parallel to **ê**_2_), and the (*x*_1_, *s*) plane coincides with a principal axis of the cross section; (iii) the elastic deformations are small (*ε*_e_ ≪ 1); and (iv) as before, the cross section of the entire bundle is planar and constant. These assumption simplify the elastic energy of the organ per unit arc length, which is given by the integral 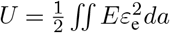 on the cross section. Expressing the elastic strain of each concentric shell *ε*_e,*i*_ using Eq. 2 gives:

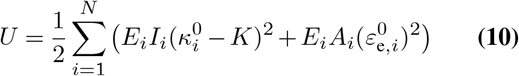

where *E*_*i*_ is the effective Young’s modulus of shell *i, I*_*i*_ is its second moment of area about the axis normal to the plane of bending, *A*_*i*_ is the area of its cross section, and *K* is the actual curvature of the bundle which acts as the actual curvature of each concentric shell. Since the dynamics are quasi-static, as the shells remodel their intrinsic lengths and curvatures, the energy landscape evolves and the system remains at its instantaneous minimum.

#### Curvature dynamics

To express the shape dynamics, we begin with the rod’s curvature. Minimizing the energy *U* (Eq. 10) by *K* gives the intrinsic curvature *K*^0^:

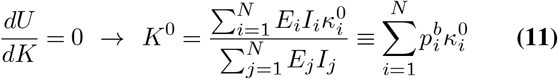

where in the last equality we define:

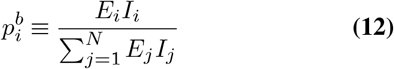

as the elastic bending weight of shell *i*, such that 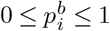 and 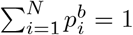. In equilibrium, in the absence of external forces, the bundle assumes the curvature *K*^0^ such that *K* = *K*^0^. Residual bending stresses then appear as long as the intrinsic curvatures of individual shells 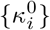 are different from *K*^0^. Following Eq. 6, the dynamics of intrinsic curvature of each shell are given by:

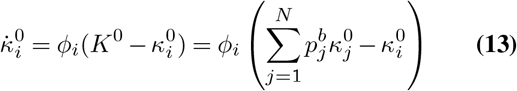

where in the second equality we substituted Eq. 11. We solve these dynamics after introducing the dynamics of the average strain rate.

#### Axial growth dynamics

The dynamics of the average strain rate and the average elastic strain in each shell are given by Eq. 5:

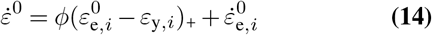

where the average strain rate 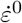 is independent of *i* and assumed uniform across each cross section, keeping the cross sections planar. To continue, we now express 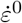 using an averaged sum of the growth rates of the individual shells. For this, we first note that in equilibrium the sum of residual stresses within the organ vanishes, which from symmetry can be applied to each cross section via:

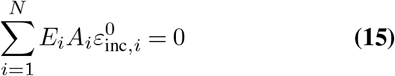

where 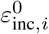 is the average incompatible strain in shell *i*. We then express the hydrostatic elastic strain, which is constant across the section, using:

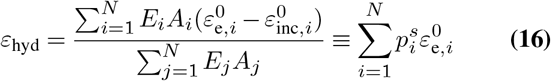

where in the first equation we substitute Eq. 9, and in the second equation we use Eq.15 and define:

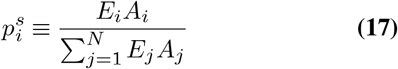

as the elastic stretch weight of shell *i*, such that 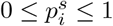 and 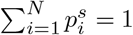. Multiplying Eq. 14 by 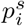 and summing over the *N* shells gives the actual strain rate:

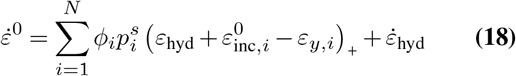

where the sum of the time derivatives of the residual axial elastic strains vanishes following Eq. 15. For brevity, we denote the elastic strain that drives growth by:

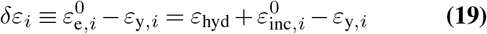

such that the growth law in Eq. 4 can be written as 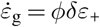. Using this notation, substituting Eq. 18 in Eq. 14 allows to write the dynamics of the axial elastic strain of shell *i* as:

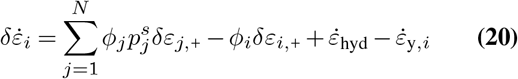

#### Solving the dynamics

The dynamics of curvature and average axial strain rate of the concentric shells depicted in Eqs. 13,20 can each be written as two sets of N coupled ODEs. Denoting the vectors 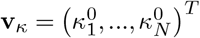 and **v**_*δε*_ = (*δε*_1,+_, …, *δε*_*N*,+_)^*T*^ allows us to write Eqs. 13,20 as:

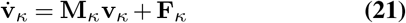

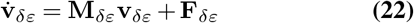

where **M**_*κ*_ and **M**_*δε*_ are *N* × *N* matrices defined by the corresponding elastic weights (Eqs. 12,17) and shell extensibilities:

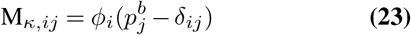

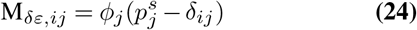

where *i, j* ∈ {1, …, *N*} and *δ*_*ij*_ is the Kronecker delta. The dynamics defined by these matrices describe the quasi-static growth response to incompatibility-induced elastic strains, such that the steady-state values of curvature and axial elastic strain lie in the kernel of these matrices. The vectors **F**_*κ*_ and **F**_*δε*_, which render Eqs. 21,22 inhomogeneous, represent processes other than incompatibility that varies the steady state, such as time-dependent growth parameters, temporal variations in turgor pressure, or transverse gradients in growth parameters within a single shell, as shown in Eqs. 8,20. For example:

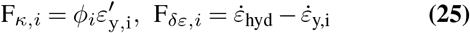

The general solution to Eqs. 21,22 with constant matrices **M**_*κ*_ and **M**_*δε*_ can be written as (46) (removing the subscripts for clarity):

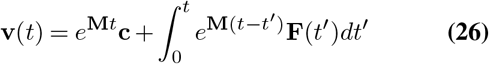

where **c** is a vector of constant coefficients set by the initial conditions. Using the solutions given by Eq. 26, the elastic strain profile within each shell can be written using Eq. 2. We note that the dynamics become more complex when stiffness profiles and extensibilities are allowed to vary in time (see SI).

### Coupling the epidermis and inner tissues (N = 2)

To demonstrate the richness of this analytically tractable model and explore its predictions, we focus on the simplest case of two shells, which may represent the epidermis and inner tissue in a rod-like organ. This effectively models the “epidermal growth control” hypothesis, and allows us to explore the consequences and observables when we assume that the epidermis mechanically controls the organ’s shape dynamics. Without loss of generality, we identify shell 1 with the inner tissues (denoted by *in*) and shell 2 with the epidermis (denoted by *ep*). For simplicity, in all of the following examples we assume **M**_*κ*_ and **M**_*δε*_ are time independent and use the solution given by Eq. 26 (the relevant matrix exponentials *e*^**M***t*^ for N=2 appear in the SI).

#### Stress relaxation with inhomogeneous growth and elastic parameters

We begin by assuming the dynamics are set only by incompatibility-induced elastic strains, such that **F**_*κ*_ = 0 and **F**_*δε*_ = 0. In axial growth dynamics, assuming all tissues actively grow with *δε*_*i*_(*t*) *>* 0, the elastic strains decay exponentially to steady state values:

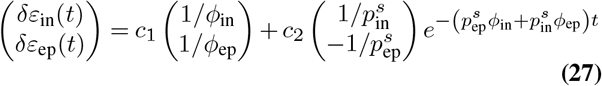

where *c*_1_ and *c*_2_ are integration constants set by the initial conditions. The resulting strain rate of the organ is then given by Eq. 18:

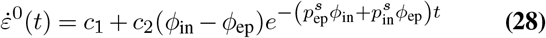

such that in this edge case temporal variations occur only if *ϕ*_in_ ≠ *ϕ*_ep_. For example, setting *δε*_in_(0) = *δε*_ep_(0) = *ε*_hyd_ – *ε*_y_ and *ε*_inc,1_(0) = *ε*_inc,2_(0) = 0 we obtain:

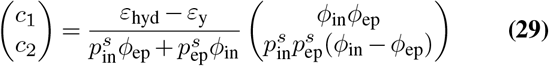

We validate the steady state of this model against a model of a continuous, straight, inhomogeneous rod and find good agreement (see SI). From a biological perspective, Eqs. 27,28 could be used to investigate a scenario in which external signals, such as light or heat, trigger an abrupt change in the axial strain rate of an organ, as it both describes the dynamics as well as give predictions regarding the distribution of residual stresses.

Addressing the yield threshold by letting *δε*_*i*_(*t*) ≤ 0 introduces a nonlinear component into the dynamics which, as discussed in a later example, enables prediction of growth-zone lengths via temporal growth cessation. Interestingly, it also predicts non-trivial elastic relaxation dynamics following the insertion of new axial cell walls during division, as we discuss now. For this, we label the new walls as *new* and the preexisting tissue as *old* (see Fig. 3). Since adding axial walls increases the effective cross-sectional stiffness ⟨*E*⟩, we set 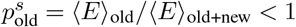. We assume the new walls initially carries no mechanical load, so that 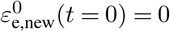 and *δε*_new_(*t* = 0) = − *ε*_*y*_ *<* 0, while the remaining tissue has *δε*_old_(*t* = 0) = *P/* ⟨*E*⟩ _old_ − *ε*_*y*_ *>* 0. The dynamics then describe how coupling to the growing tissue increases the elastic stretch of new walls until *δε*_new_ = 0, at which point the Heaviside nonlinearity activates and the new walls begins to grow. At the same time, the preexisting walls are held back by the new walls, lowering their elastic strain and the observed strain rate. For *ϕ*_new_ = *ϕ*_old_ ≡ *ϕ*, activation of the new walls occurs only if *P/* ⟨*E*⟩ _old+new_ *> ε*_*y*_, and the system then relaxes exponentially with rate *ϕ* to the reduced steady growth rate *ϕ* (*P/* ⟨*E*⟩ _old+new_ − *ε*_*y*_). This example demonstrates how discrete divisions can alter both the local mechanical landscape and growth dynamics.

**Fig. 3.**
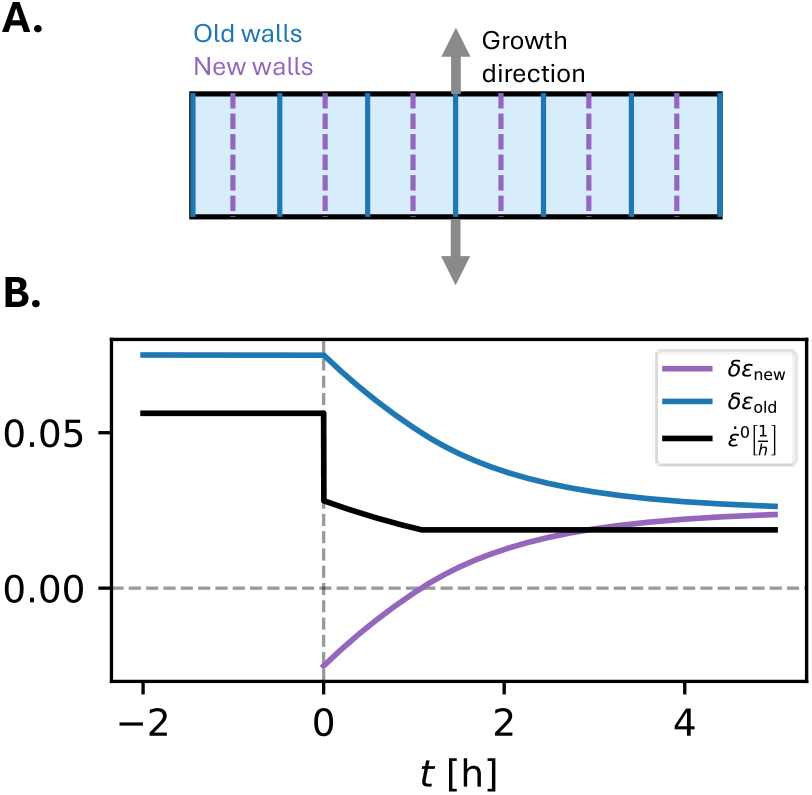
Mechanical effect of axial division. A. Model illustration: At time *t* = 0 cells divide longitudinally, adding new cell walls which are unstretched and tug on the preexisting growing cell walls. **B**. The dynamics of the visible axial strain rate and the growth-inducing part of the elastic stretches *δε* of both the new and old walls. Initially, the new cells walls do not actively grow, and passively hinder the ability of the growing cell walls to extend the tissue, while their elastic stretch increases. When their elastic stretch equals the yield threshold (*δε*_old_ = 0), the new walls start to actively grow, leading to a new steady growth rate which is lower than the initial rate assuming constant turgor. The parameter values used for this example are: *ϕ* = 0.75, 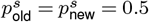 , *ε*_y_ = 0.025, *δε*_new_(*t* = 0) = 0.075. In the case of a single division, the elastic stretch weight of the new wall 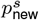 might be very small, decreasing the magnitude of the predicted discontinuity in the dynamics.

For completeness, we note that the solution of curvature dynamics also presents an exponential decay to steady state values, and the dynamics of the apparent rod’s curvature are found by Eq. 11:

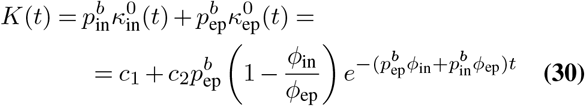

where, similar to Eq. 28, *c*_1_ and *c*_2_ are integration constants set by the initial conditions and relaxation only occurs if *ϕ*_in_ ≠ *ϕ*_ep_. Given the initial values of the intrinsic curvatures of the shells 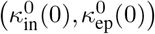 yields:

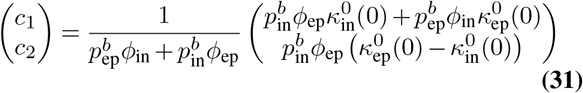

Overall, these examples show that tissue heterogeneity generates incompatibility-induced elastic strains. In turn, this leads to exponential relaxation of axial stress toward steady-state values determined by the initial conditions, extensibilities, and elastic weights. We now turn to examples of more complex shape dynamics that arise when growth or elastic parameters vary continuously in time due to aging or genetic regulation.

#### Epidermal control of curvature leads to mechanical “memory” and effective autotropism

During tropic movements, plants regulate their curvature for posture control and in response to external directional signals. To demonstrate how residual stresses might affect these growth driven turns, we now model an epidermal control of curvature. Specifically, we assume the epidermis reacts to a morphogen which does not affect the inner tissues, such that F_*κ*,ep_(*t*) = *f* (*t*) ≠ 0 and F_*κ*,in_ = 0. For example, this might represent the response to anisotropic auxin patterns during gravitropic movements, which in roots is modeled via *f* (*t*) ∝ sin(*θ*_*tip*_(*t*)), where *θ*_*tip*_(*t*) is the inclination of the root apex with respect to gravity (20). In this case, as the epidermis varies its intrinsic curvature, it bends the inner tissues, which relax to their current curvature with rate *ϕ*_in_ (as in Eq. 6). The inner tissues therefore “lag” behind the epidermis in their intrinsic curvature, resulting in dynamic curvature incompatibility. Assuming the initial organ is straight with *K*(0) = 0, the resulting dynamics of the curvature of the rod can be written as:

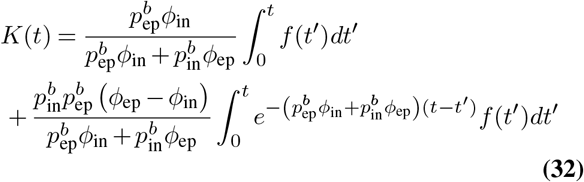

An example of this model is shown in Fig. 4. To analyze the resulting dynamics, we note that time derivative of Eq. 32 gives:

**Fig. 4.**
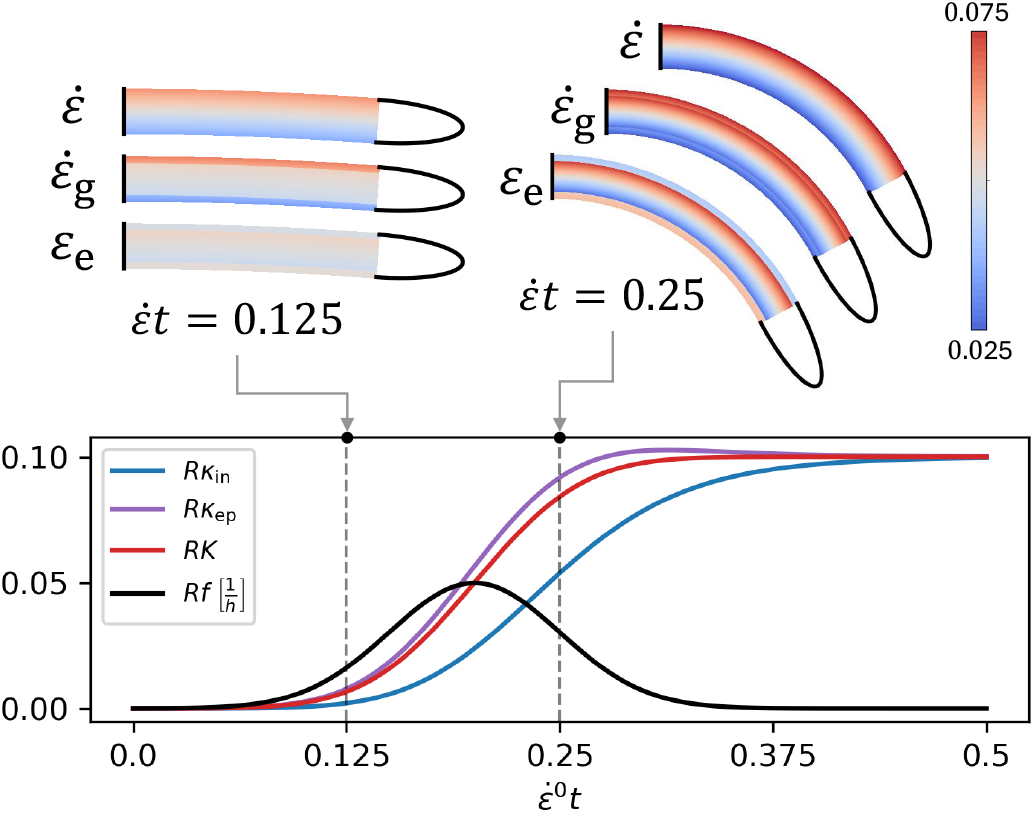
Epidermal control of curvature. Bottom: The active curvature rate of the epidermis *f*(*t*) leads to time dependent intrinsic curvatures which are incompatible with the actual curvature. These are plotted against the non dimensional time 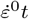 where 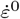 is the constant axial growth rate of the centerline. The vertical dashed lines mark the snapshots shown above, and *R* is the radius of the cross section. Top: Snapshots of the dynamics, portraying the actual strain rate, growth rate and axial elastic strain of a bending organ. At 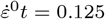, the curvature rate of the epidermis is shown and drives the curvature dynamics. At 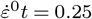, the elastic strain shows that due to incompatibility the inner tissues are elastically bent and the epidermis is elastically straightened, affecting the overall dynamics. The parameter values used for this example are: *ϕ*_in_ = *ϕ*_ep_ = 1 1/hour, *ε*_hyd_ = 0.05, 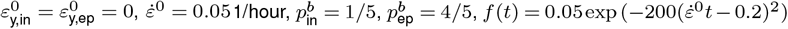.

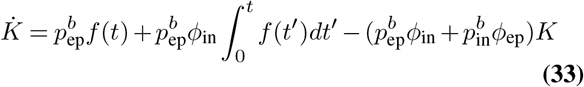

The three additive terms in the right hand side of Eq. 33 can be interpreted as: (i) an immediate effect of the curvature rate of the epidermis, (ii) mechanical “memory” coming from the relaxation rate of the inner tissue (21), and (iii) effective autotropism (19, 20, 47). Autotropism is a growth induced straightening motion which acts alongside gravitropic responses in tilted shoots and roots. Though autotropism is often linked to active proprioception of the organ’s own curvature, its underlying mechanisms remain unclear, and a specific morphogen-mediated curvature sensing pathway is yet to be found. The model in Eq. 33 raises the possibility that autotropism may be a relatively passive process, resulting from the joint relaxation of bending residual stresses within the organ, with sensitivity:

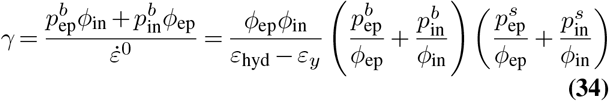

where in the second equality we use, for example, the steady-state strain rate given by Eqs. 28,29. For instance, if *ϕ*_in_ = *ϕ*_ep_ we obtain *γ* = 1*/*(*ε*_hyd_ − *ε*_*y*_), which, assuming *ε*_hyd_ − *ε*_*y*_ ∼ 0.1, gives an order-of-magnitude estimate *γ* ∼ 10. As we show in the SI, when *ϕ*_in_ ≠ *ϕ*_ep_ the allowed range of *γ* spans several orders of magnitude, including the experimentally inferred value *γ* ∼ 1 reported for roots (20). Possible supporting evidence to this model appear in (48), where peeling of plant shoots following a gravitropic response revelead asymmetric stress distribution across the bent cross-section, similar to those illustrated in Fig. 1D and Fig. 4A.

To continue the analysis, we note that the steady-state curvature from Eq. 33 in the limit *t* → ∞, assuming *f* (*t* → ∞) = 0, can be written as:

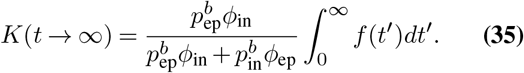

This expression shows that once the active bending of the epidermis encoded by *f* (*t*) has ceased, and as long as growth-induced remodeling continues, the steady-state curvature is determined by the time integral of the epidermal curvature rate. This implies that although the model for dynamic autotropism in Eq. 34 is general, achieving straight segments at steady state requires the right-hand side of Eq. 35 to be close to zero. For example, this occurs when 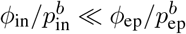, corresponding to a regime in which the inner tissues remain effectively intrinsically straight throughout the dynamics and act as a passive bending spring. A similar straightening mechanism was considered in (49), where the intrinsic angles between cell walls were constrained to right angles. In addition, this limiting case may reflect the role of cellular microfibrils in posture control observed experimentally in (50).

#### Self-similar axial growth profile driven by a varying yield threshold

So far, we have used our framework to analyze growth and elastic inhomogeneities between tissues within the same cross section, effectively working in Lagrangian coordinates that move with the material segments as they grow. In most experimental settings, however, growth patterns are phenotyped in Eulerian coordinates, with positions measured by arc length from either the apex or the base of the organ. To demonstrate how our framework can be applied to such descriptions, we now consider the self-similar triangular axial growth profile observed in roots (42, 51). As we show below, the underlying self-similar symmetry allows the growth dynamics to be formulated using a stationary Eulerian description.

We begin by assuming for simplicity that *ϕ*_ep_ = *ϕ*_in_ ≡ *ϕ* and that the dynamics are driven by a time dependent yield threshold of the inner tissues, such that 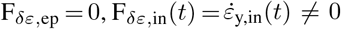 and 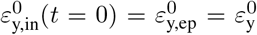. The temporal changes in the yield threshold are assumed to be regulated by a spatial morphogen pattern, linking the mechanical response of the tissue to an underlying biochemical patterning. Furthermore, as we show in the following, regulating the growth of the inner tissues causes the epidermis to mechanically restrict and mediate growth, in agreement with the “epidermal growth control” hypothesis. Concretely, these assumptions give:

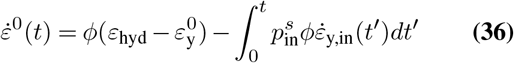

and:

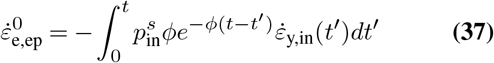

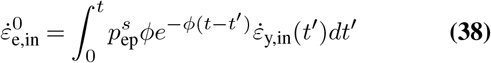

such that the elastic strain rate in both the inner tissues and the epidermis results from a convolution between the varying yield threshold 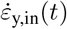 and an effective memory kernel with timescale 1*/ϕ*.

To enable comparison with experimental measurements, we now assume that both the morphogen pattern driving the dynamics as well as the resulting axial growth rate are self-similar, or stationary with respect to the apex of the organ. We denote the arc-length from the apex via *s*, such that 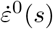. Mathematically, this gives a stationary velocity 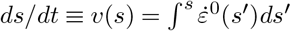,which describes the axial displacement of material points (18). The corresponding characteristic curves then satisfy:

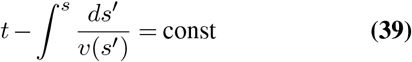

Because the motion is unidirectional and stationary, a material point on the epidermis located at a given distance *s*_0_ *>* 0 from the apex at time *T* moves according to a time-invariant characteristic. Consequently, after a fixed time interval *δt*, it reaches a uniquely determined downstream position *s*(*T* + *δt*) (see Fig. 5A). This follows from the time-translation invariance of Eq. 39. Without loss of generality, we now focus on a specific characteristic associated with a given *s*_0_ *>* 0, such that *t*(*s*_0_) = 0. With a slight abuse of notation, we denote this characteristic by the mapping *s* = *s*(*t*). This mapping can be obtained directly from the solution for the relative growth rate 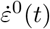 derived from our concentric shells model (Eq. 36). To show this, we begin by noticing that the strain rate can be related to the velocity via (18):

**Fig. 5.**
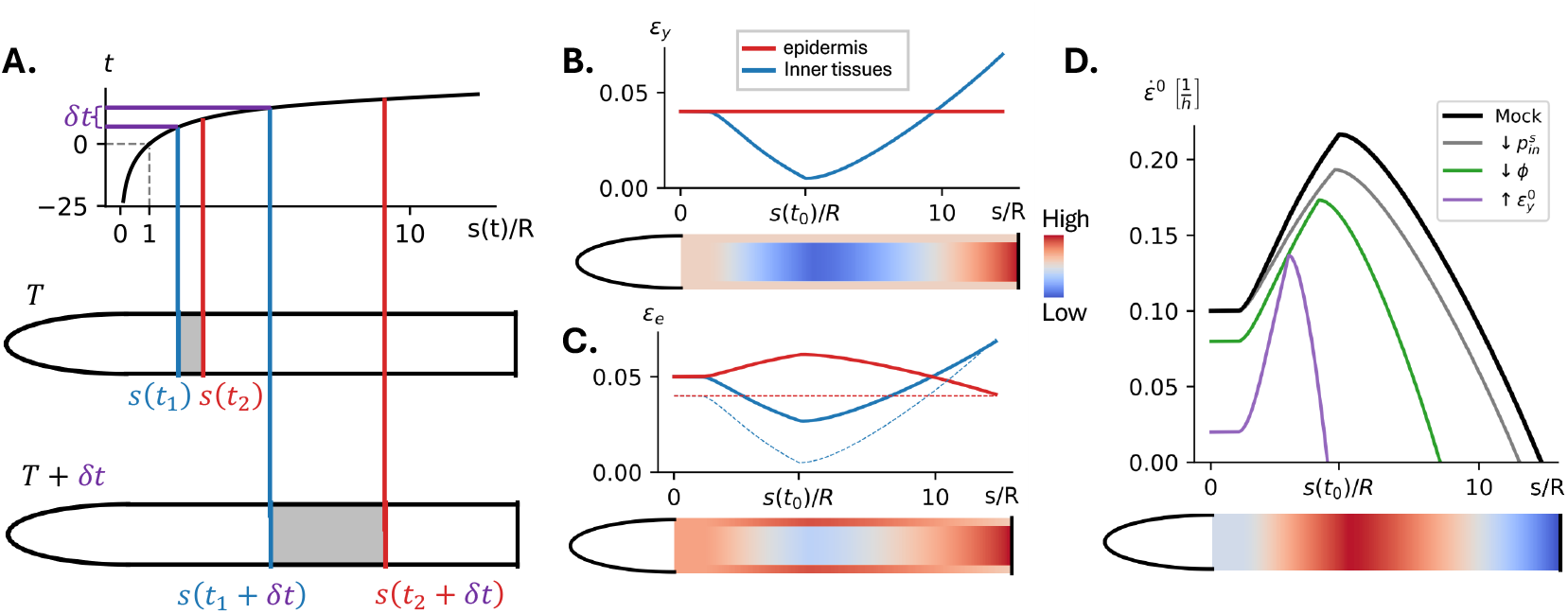
A. Self-similar axial growth illustration. Top: as the whole organ expands symplastically, material points move along the organ length, changing their axial position (*s*) over time (*t*) along characteristic trajectories. In the case of self-similar growth, the motion of each material point can be described by a single trajectory *s*(*t*), determined from the axial strain rate 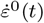. Here, we set *s*(*t*) such that *s*(0) = *R*. Since the apex is a fixed point (*v*(*s* = 0) = 0), approaching *s →* 0_+_ corresponds to *t → −∞* along the characteristic. Bottom: motion of two material points along the organ axis as described by *s*(*t*). At any time *T* , the points are located at *s*_1_ = *s*(*t*_1_) and *s*_2_ = *s*(*t*_2_), where *t*_1_, *t*_2_ denote the times elapsed since the material points passed *s* = *R*. After an additional time interval *δt*, their new locations are *s*_1_ = *s*(*t*_1_ + *δt*) and *s*_2_ = *s*(*t*_2_ + *δt*). **B-D. Self-similar dynamics of axial growth and elastic strain due to a differential yield threshold profile**. The model is solved in time and plotted against arc-length via *s*(*t*), normalized by the cross section radius *R*. Below each graph, the data is also presented as a heatmap on an illustration of the organ separated into epidermal and inner tissues. The parameter values used for this example are: *ϕ* = 10 1/hour, 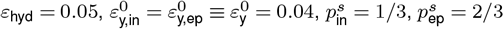, *a* = 0.05, *b* = 0.5, *t*_0_ = 14 hours. **B**. The given yield threshold which drives the dynamics, constant in the epidermis and varying in the inner tissues. **C**. The resulting profile of axial elastic strain (solid lines). The yield thresholds appears in dashed lines for comparison. We note that *δε* which drives growth appears as the difference between the solid and dashed lines. **D**. The profile of actual strain rate given by the mode mimics the strain rate profile observed in roots (“Mock”). For comparison, we plot the stationary strain rates resulting for perturbation of various model parameters by 20 percent (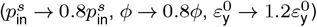), where *s*(*t*) is recalculated for each graph based on the resulting 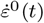.

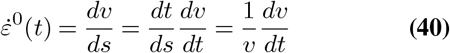

where in the second equality we used the chain rule. Next, we use Eq. 40 to express the velocity as a function of time via 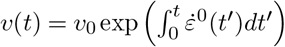 , where *v*_0_ is the velocity at *s*_0_ (or *t* = 0). Integrating this expression in time gives the space-time bijection following 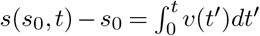.

To use this symmetry and reproduce the triangular growth profile found in roots, we now assume the yield threshold rate is a skewed triangular function of time:

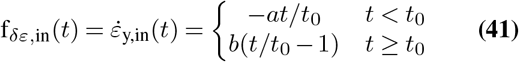

with 0 *< a < b*, which allows to solve the dynamics analytically. As an example, we assume *s*_0_ = *s*(*t* = 0) = *R* and that the growth rate between 0 ≤ *s* ≤ *R* is constant and equals 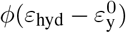. The results are plotted in Fig. 5B-D against arc-length. Initially, the lowering of the yield threshold of the inner tissue (Fig. 5B) drives the strain rate acceleration (Fig. 5D, ‘Mock’), and causes the epidermis (inner tissues) to accumulate relative stretch (compression), which is maintained for most of the dynamics (Fig. 5C). As the yield threshold of the inner tissues begins to increase, the signs of the integrals in Eqs. 37,38 are reversed, and the elastic stretch of the epidermis begins to decrease. At the point where growth ceases at the base, the epidermis is compressed with respect to the inner tissues over a short arc length (Fig. 5C).

Such relative compression may lead to a twist instability, which appears in mutant phenotypes between the growth zone and mature zone of roots (25, 52). Lastly, in Fig. 5D we demonstrate the model predictions regarding changes in the axial strain-rate profile and growth zone length following variations in elastic parameters 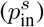 and growth parameters 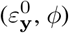. These variation may arise, for example, from the exogenous applications of a growth-regulating morphogen that induces mechano-chemical activity. In the present formulation, the perturbations obey the time varying yield described by Eq. 41. In future work, however, spatial variations in growth and elastic parameters will be prescribed by a super-imposed diffusing morphogen field.

## Conclusion

We introduce a mathematical framework that couples growth and elasticity in rod-like plant organs, enabling systematic investigation of the interplay between incompatibility-induced residual stresses and macroscopic morphogenesis. By representing organs as connected morphoelastic cylindrical shells, the governing equations reduce to a tractable form that enables quantitative predictions of shape and elastic strain dynamics in the presence of tissue heterogeneity. This framework provides direct insight into how growth couples to internal mechanical loading, and suggests that residual stresses are not merely a by-product of growth but can play a key role in driving plant morphogenesis. Specifically, using a simplified two-compartment model inspired by the “epidermal growth control” hypothesis, we find that: (i) autotropism may arise from relaxation of residual stresses during bending motions; (ii) structural incompatibilities can lead to a form of mechanical “memory”; and (iii) cell divisions can modify local stiffness profiles and tissue-scale growth dynamics. Most of the conceptual mechanisms suggested by our framework require mechanical testing to validate or falsify, particularly since growth patterns may remain mechanically compatible (as in Eq. 8). The residual stress patterns predicted by our framework can be probed using laser ablation, as well as peeling and splitting assays (1, 48, 53, 54). Finally, although we focus on rod-like organs, similar theoretical approaches could shed light on the role of residual stresses in shaping morphodynamics in systems with different geometries, including planar leaves and dome-shaped apical meristems.

## Acknowledgments

A.P. acknowledges support from the Gatsby Charitable Foundation (grant GAT3395/PR4B) and the Herchel Smith Fund, and thanks Yasmine Meroz, Anne-Lise Routier-Kierzkowska, Henrik Jönsson, Bruno Moulia, Mathieu Rivière and Sarah Robinson for the insightful discussions.

## Supplementary Information

### Morphoelastic rods: state-of-the-art

#### Shape dynamics

The position vector of each point in the rod

***p*** can be written as (see Fig. 1B):

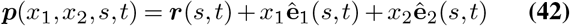

where the cross sections are bound to move together as solid planes, and the tangent to the centerline 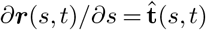 acts as their normal. Assuming a shearless and twistless rod, the rotation of the cross sections along the organ is captured by the curvature or Darboux vector of the centerline 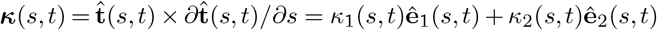. To let the rod grow axially, we use the arc-length of the rod at time *t* = 0 denote by *S*_0_ as a material coordinate, let the actual arc length develop in time via *s* = *s*(*S*_0_, *t*) and define the axial stretch as *λ*(*s, t*) = *∂s/∂S*_0_. Using 42, the metric in the rod with respect to the coordinate system (*x*_1_, *x*_2_, *S*_0_) is then (omitting the dependencies for brevity):

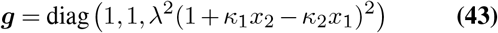

Under these constraints, the material frame acts as a Bishop (rotation-minimising) frame, and the deformation rate tensor can then be found by taking the co-rotational time derivative of the metric from Eq. 43 (55). Denoting 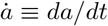, this gives in the orthonormal basis (16):

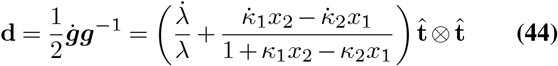

Using the logarithmic strain measure *ε*^0^ = ln(*λ*) and assuming small curvature *R* |***κ***| ≪ 1, where *R* is the maximal transverse extent of the cross section, gives Eq. 1 from the main text.

#### Rod mechanics

Separation of timescales between slow growth and fast elastic relaxations allows us to assume quasi static growth (17). Mechanical equilibrium is then expressed by the time independent Cosserat rod equations along the centerline (31, 56):

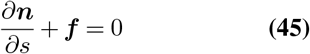

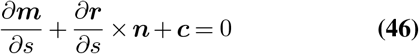

where ***n*** are the internal contact forces, ***m*** are the internal torques or bending moments, ***f*** are the external forces per unit length and ***c*** are the external couples per unit length, all functions of *s* and *t*. Since in our model the cross sections remain planar, the internal contact forces and bending moments can be expressed using the 0-th and 1st moments of the axial stress (31, 57):

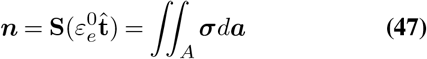

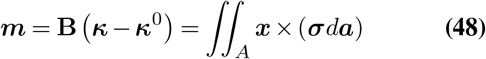

where 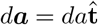 is an infinitesimal area vector on the cross section, and **S** and **B** are stiffness matrices which are diagonal in the local material coordinates, where S_33_ = ∫∫_*A*_ *Eda* ≡ ⟨*E*⟩ *A*, denoting ⟨*E*⟩ as the average axial stiffness of the cross section, and 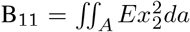 and 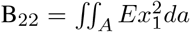 are the second moments of the axial stiffness (second moment of area).

By definition, the sum of residual stresses within the organ vanishes, which from symmetry can be applied to each cross section:

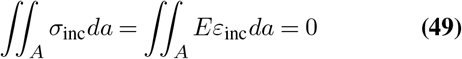

The reaction strain to the inner hydrostatic pressure can be derived from Eq. 47 assuming the turgor pressure *P* is constant:

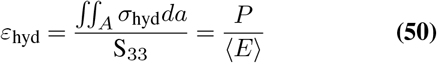

Accordingly, in case the turgor pressure maintains a transverse gradient such that *σ*_hyd_ = *P*_0_ + ***x*** · **∇** *P* , Eq. 48 gives the reaction bending strain:

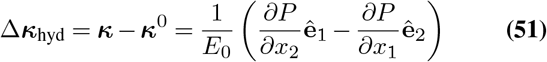

Lastly, in the case of a transvers gradient in stiffness *E* = *E*_0_ + ***x*** · **∇** *E* and constant turgor, Eq. 48 gives a different reaction bending strain:

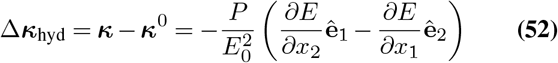

Eqs. 51,52 give Eq. 7 in the main text.

#### Intrinsic curvature dynamics

To derive the dynamics of intrinsic curvature, we note that by assuming a constant cross section and stiffness profile, the co-rotational time derivative of Eq. 48 gives:

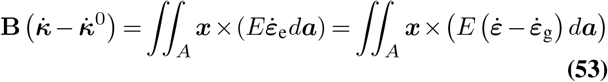

where we used Eq. 3 in the first equality and Eq. 5 in the last equality. Substituting the strain rate from Eq. 1 in Eq. 53 then gives the dynamics of intrinsic curvature (see also (9)):

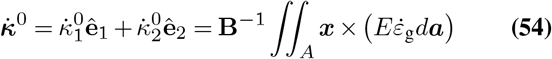

Substituting the strain-based growth law from Eq. 5 in Eq. 54 while assuming *ε*_e_ *> ε*_y_ gives:

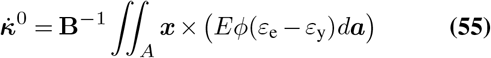

In the case of homogeneous growth parameters, substituting Eq. 48 in Eq. 55 yields Eq. 6 in the main text. Adding the inhomogeneities 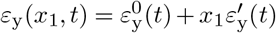 and *ϕ*(*x*_1_, *t*) = *ϕ*^0^(*t*)+*x*_1_*ϕ*^*′*^(*t*) while keeping the transverse gradients to first order results in Eq. 8.

#### Residual stress in a straight inhomogeneous rod

Here, we investigate the steady state of a free, straight rod growing axially with a constant strain rate 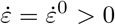. To avoid changes in the intrinsic curvature of the rod, we assume the elastic and growth parameters of the rod are axis symmetric, such that the rod is inhomogeneous only with respect to the radial distance form the axis *x* = |***x***| . In addition, we assume the elastic strain profile of the rod is constant yet not necessarily homogeneous with 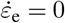. The elastic strain can then be expressed by inverting Eq. 5:

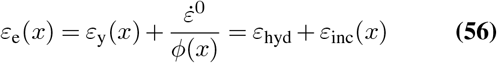

where the last equality comes from Eq. 9. Equation 56 shows that differential yield strain *ε*_y_ or extensibility *ϕ* vary the local steady-state elastic strain and corresponding intrinsic length. The possible profiles of these growth parameters are constrained by the condition for mechanical equilibrium, which can be found by integrating Eq. 56 multiplied by the local stiffness *E*(*x*) across the cross section and substituting Eqs. 49-50:

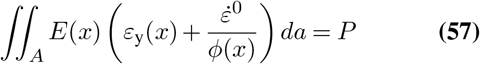

This model agrees with the steady state predicted by Eqs. 27-29 in a piecewise manner: substituting the steady state growth rate *c*_1_ from Eq. 29 in Eq. 27 reproduces Eq. 56, as well as Eq. 57 via 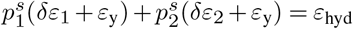.

We note that this model allows to describe the local relative residual stress, which we define by the difference between the axial stress to its average value *P* (Eq. 57):

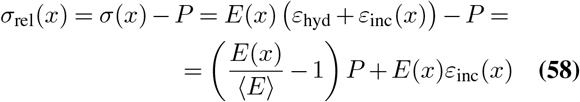

where in the last equality we used Eq. 50. Equation 58 shows that the relative stress can result from two different additive mechanisms (3): (i) turgor pressure, which distributes stress unevenly as in parallel springs: if the local stiffness is higher (smaller) than the average stiffness, the tissue is relatively taut (compressed); and (ii) growth induced length incompatibility, which as shown Eq. 56 can be attributed to differential growth parameters.

#### Matrix exponentials used in the main text

In the case *N* = 2, the matrices given by Eqs. 23,24 give:

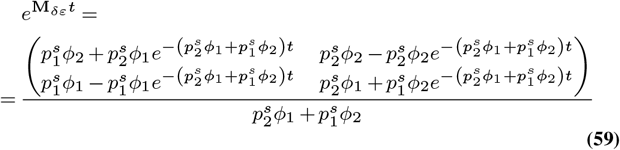

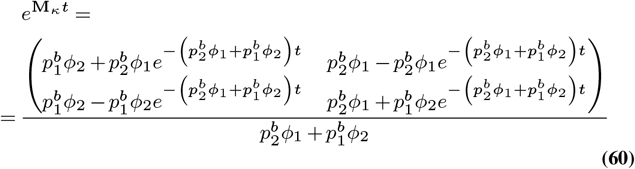

#### Note on Time-Dependent Stiffness

Although we do not solve the dynamics for time-dependent stiffness profiles 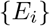, we briefly note the complications such profiles introduce. First, both the elastic weights 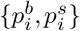 and the hydrostatic strain *ε*_hyd_ = *P/* ⟨*E*⟩ become time-dependent. Second, when deriving the axial strain dynamics, taking the time derivative of Eq. 15 produces two sums. This modifies Eq. 18 by adding the following extra term on the right-hand side:

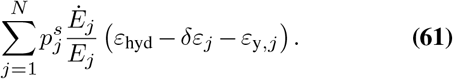

As a result, the updated matrices **M** becomes time-dependent, and the system can be solved using the fundamental matrix method (46). The same applies if the extensibilities {*ϕ*_*i*_} are time-dependent.

#### Estimation of the autotropic sensitivity *γ*

Here, we estimate the order of magnitude for *γ* predicted by our model in Eq. 34, by assuming *ε*_hyd_ − *ε*_*y*_ = 0.1. To match the bending and stretching elastic weights in a realistic manner, we assume the cross section of the inner tissues is cricular with radius *R* and that the epidermis is a thin layer around the inner tissues of width *t*. This allows to calculate their areas and second moment of inertia explicitly, such that 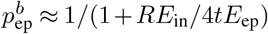 and 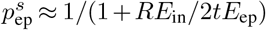. The orders of magnitude of *γ* are then portrayed in Fig. 6. We note that different models for the average axial strain rate 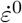 will lead to different functional forms for *γ*, varying its possible values.

**Fig. 6.**
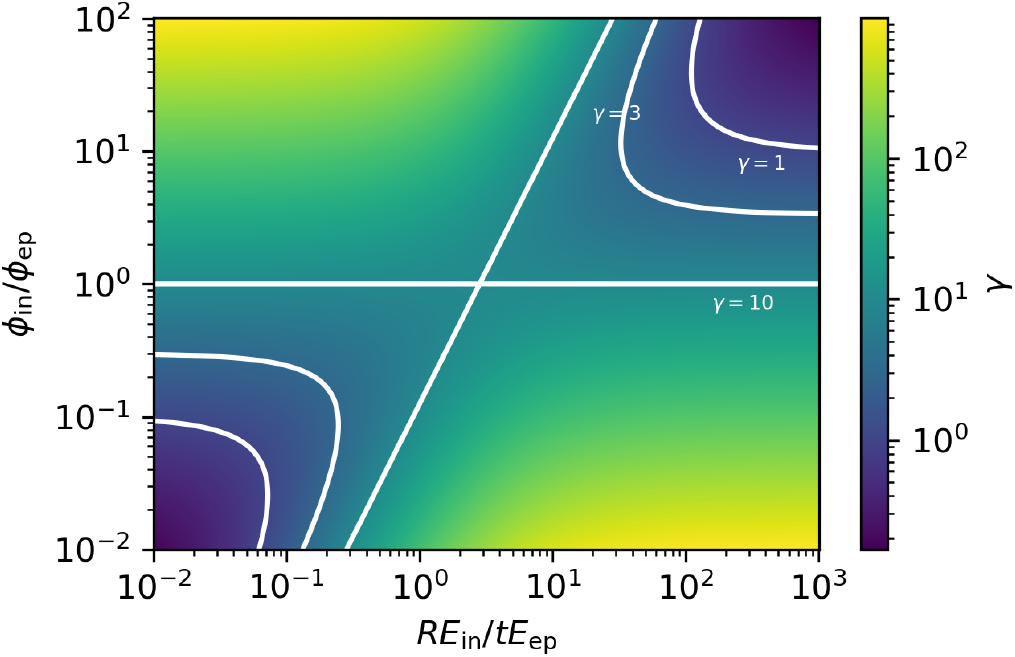
**Estimation of the autotropic sensitivity** *γ* based on Eq. 34. Here, we use *ε*_hyd_ *− ε*_*y*_ = 0.1, 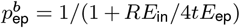 and 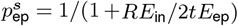.

**Table 1.** Model parameters and variables used throughout the manuscript.

| Symbol | Description | Units |
| --- | --- | --- |
| $r(s, t)$ | Centerline position vector of the organ | length |
| $s$ | Arc-length coordinate along the organ | length |
| $t$ | Time | time |
| $\hat{e}_1, \hat{e}_2, \hat{e}_3$ | Local orthonormal material frame | — |
| $\hat{t}$ | Tangent vector to the centerline ($\hat{t} = \partial r / \partial s$) | — |
| $x = (x_1, x_2)$ | Coordinates on the cross section | length |
| $R$ | Cross section radius | length |
| $\kappa = (\kappa_1, \kappa_2)$ | Actual curvature vector | length<sup>-1</sup> |
| $\kappa^0 = (\kappa_1^0, \kappa_2^0)$ | Intrinsic curvature vector | length<sup>-1</sup> |
| $K$ | Actual curvature of the shell bundle | length<sup>-1</sup> |
| $K^0$ | Intrinsic curvature of the shell bundle | length<sup>-1</sup> |
| $\dot{\varepsilon}$ | Axial strain rate | time<sup>-1</sup> |
| $\dot{\varepsilon}_0$ | Midline axial strain rate | time<sup>-1</sup> |
| $\varepsilon_e$ | Elastic axial strain | — |
| $\varepsilon_e^0$ | Average elastic axial strain within a shell | — |
| $\varepsilon_{\text{hyd}}$ | Hydrostatic elastic strain induced by turgor | — |
| $\varepsilon_{\text{inc}}$ | Residual elastic strain due to incompatibility | — |
| $\varepsilon_y$ | Yield strain threshold | — |
| $\delta\varepsilon$ | Growth-driving elastic strain, $\delta\varepsilon = \varepsilon_e^0 - \varepsilon_y$ | — |
| $(x)_+ \equiv x\Theta(x)$ | activation function, defined by $x\Theta(x) = x$ for $x > 0$ and $x\Theta(x) = 0$ for $x \leq 0$ | — |
| $\dot{\varepsilon}_g$ | Growth strain rate | time<sup>-1</sup> |
| $\phi$ | Effective extensibility | time<sup>-1</sup> |
| $E$ | Effective Young's modulus | pressure |
| $P$ | Internal hydrostatic turgor pressure | pressure |
| $A_i$ | Cross-sectional area of shell $i$ | length<sup>2</sup> |
| $I_i$ | Second moment of area of shell $i$ | length<sup>4</sup> |
| $p_i^s$ | Elastic stretch weight of shell $i$ | — |
| $p_i^b$ | Elastic bending weight of shell $i$ | — |
| $N$ | Number of concentric shells | — |
| $f(t)$ | Biochemically imposed curvature-rate input | length<sup>-1</sup><br>time<sup>-1</sup> |
| $\gamma$ | Effective autotropic sensitivity | — |
| $v(s)$ | Axial material velocity | length time<sup>-1</sup> |

